# Unique molecular markers for GC-D-expressing olfactory sensory neurons and chemosensory neurons of the Grueneberg ganglion

**DOI:** 10.1101/346502

**Authors:** Zhi Huang, Arthur D. Zimmerman, Steven D. Munger

**Affiliations:** Center for Smell and Taste; Department of Pharmacology & Therapeutics; Department of Medicine, Division of Endocrinology, Diabetes and Metabolism, University of Florida, Gainesville, FL, 32610, USA

**Keywords:** olfaction, phosphodiesterase, CART, guanylyl cyclase, olfactory bulb, glomeruli

## Abstract

The main olfactory bulb (MOB) is differentiated into subregions based on their innervation by molecularly distinct chemosensory neurons. For example, olfactory sensory neurons (OSNs) that employ a cGMP-mediated transduction cascade – guanylyl-cyclase D-expressing (GC-D+) OSNs of the main olfactory epithelium (MOE) and chemosensory neurons of the Grueneberg ganglion (GGNs) – project to distinct groups of “necklace” glomeruli encircling the caudal MOB. To better understand the unique functionality and neural circuitry of the necklace glomeruli and their associated sensory neurons, we sought to identify additional molecular markers that would differentiate GC-D+ OSNs and GGNs as well as their target glomeruli. We found in mouse that GC-D+ OSNs, but not other MOE OSNs or GGNs, express the neuropeptide CART (cocaine- and amphetamine-regulated transcript). Both GC-D+ OSNs and GGNs, but not other MOE OSNs, express the Ca^2+^/calmodulin-dependent phosphodiesterase Pde1a, which is immunolocalized throughout the dendrites, somata and axons of these neurons. Stronger Pde1a immunolabeling in necklace glomeruli innervated by GGNs than in those innervated by GC-D+ OSNs suggests either greater Pde1a expression in individual GGNs than in GC-D+ OSNs or a difference in sensory neuron innervation density between the two types of necklace glomeruli. Together, the unique molecular signatures of GC-D+ OSNs, GGNs and their MOB targets offer important tools for understanding the processing of chemosensory information by olfactory subsystems associated with the necklace glomeruli.

## INTRODUCTION

Mammals typically employ several distinct types of olfactory sensory neurons (OSNs) to detect the diversity of odorants in their environment (Munger et al., 2010). These different OSN subpopulations vary in the chemical stimuli to which they respond as well as in the molecular mechanisms they use to detect and transduce those stimuli into a neural signal that can be transmitted to the brain. The discovery of the odorant receptor (OR) superfamily (Buck and Axel, 1991) and its members’ differential expression across the main olfactory epithelium (MOE) (Ressler et al., 1993; Vassar et al., 1993) provided essential tools for understanding the functional diversity, anatomical arrangement and coding strategies of the canonical main olfactory system. Similarly, the identification of proteins that are differentially expressed between non-canonical olfactory subsystems is essential for isolating these OSN subpopulations and their central nervous system targets for anatomical and functional studies.

Within the main olfactory system, OSNs rely on cyclic nucleotide signaling as the primary mechanism for chemosensory transduction (Munger et al., 2010). Both the odorant receptor (OR)-expressing canonical OSNs and the trace amine-associated receptor (TAAR)-expressing OSNs (Liberles and Buck, 2006) use the second messenger cyclic 3’5’-adenosine monophosphate (cAMP) to transduce olfactory stimuli (Pace et al., 1985; Zhang et al., 2013). Though they differ in the types of G protein-coupled receptors they use to detect odorants (ORs vs. TAARs), both subpopulations share the machinery needed for cAMP signaling including adenylyl cyclase (Pace et al., 1985; Bakalyar and Reed, 1990; Zhang et al., 2013) and a cAMP-sensitive cyclic nucleotide-gated channel (Nakamura and Gold, 1987; Dhallan et al., 1990; Zhang et al., 2013). Canonical OSNs may also employ cGMP signaling on a slower time course to impact long-term odor adaptation (Zufall and Leinders-Zufall, 2000).

Several other groups of OSNs appear to use cyclic 3’5’-guanosine monophosphate (cGMP) as the primary second messenger for olfactory transduction (Zufall and Munger, 2010). However, the signaling cascades employed by some of these cells are less well characterized than are those in canonical OSNs. The cell bodies of one of the these subpopulations are found clustered in the small Grueneberg ganglia at the anterior end of the nares (Fleischer et al., 2009). Many of this heterogeneous population of ciliated chemosensory cells, which differ in morphology from OSNs of the MOE, express the the receptor guanylyl cyclase GC-G, the cyclic nucleotide-gated channel subunit Cnga3, and/or the phosphodiesterase Pde2a (Fleischer et al., 2009; Liu et al., 2009; Schmid et al., 2010; Mamasuew et al., 2011).

Two subpopulations of OSNs within the MOE also rely on cGMP signaling. One, a subset of OSNs that express the ion channel Trpc2, also contains the soluble guanylyl cyclase Gucy1b2 and the phosphodiesterase Pde6d, and uses cGMP signaling to respond to decreases in environmental O_2_ (Omura and Mombaerts, 2015; Bleymehl et al., 2016). The second expresses the receptor guanylyl cyclase GC-D (hence their typical designation as GC-D+ OSNs), Cnga3 and Pde2a (Fulle et al., 1995; Juilfs et al., 1997; Meyer et al., 2000). GC-D+ OSNs use cGMP signaling to transduce social odors involved in the social transmission of food preferences (Leinders-Zufall et al., 2007; Munger et al., 2010).

GC-D+ OSNs and GGNs have similar axonal targets in the “necklace” region of the caudal main olfactory bulb (MOB) (Juilfs et al., 1997; Roppolo et al., 2006). The axons of these two distinct chemosensory neuron subpopulations terminate in distinct but closely apposed necklace glomeruli (Matsuo et al., 2012). Thus, while the two neuronal populations are physically separated in the periphery, their central nervous system targets are difficult to distinguish without specific markers that are present in the axons and their terminals. Some molecules are not only differentially expressed across different OSN subpopulations (e.g., Pde1c and Pde4a in canonical OSNs vs. Pde2a in GC-D+ OSNs and GGNs) but also show different subcellular localization (e.g., Pde1c in cilia and Pde4a throughout the dendrites, cell bodies and axons of canonical OSNs). Here, we report that the calcium/calmodulin-dependent phosphodiesterase Pde1a is differentially localized in GC-D+ OSNs and GGNs and the cocaine amphetamine-related transcript (CART) peptide is restricted to GC-D+ OSNs. These results provide important new markers for isolating these two separate populations of chemosensory neurons.

## MATERIALS AND METHODS

### Animals

All experiments were approved by the University of Florida Institutional Animal Care and Use Committee and were conducted in accordance with the Society for Neuroscience’s Policies on the Use of Animals and Humans in Neuroscience Research. Experiments were performed in homozygous *B6;129P2-Gucy2d^tm2Mom^/MomJ* mice (GCD-EGFP mice) (Walz et al., 2007), which express enhanced green fluorescent protein (EGFP) under the control of the *Gucy2d* gene (which encodes GC-D). All animals were housed in individual microisolater cages and received food and water *ad libitum*, and the line maintained by interbreeding of heterozygotes. Male and female mice (age 4-12 weeks) were used and balanced for sex and age.

### Tissue Preparation

Mice were transcardially perfused with 4% paraformaldehyde in phosphate buffered saline (PBS, pH 7.4). Brains and nasal tissues were removed and fixed overnight in 4% paraformaldehyde in PBS, followed by cryoprotection overnight in 15% and then 30% sucrose at 4°C. 18 µm coronal sections were cut at the levels of the Grueneberg ganglion (GG) (Gruneberg, 1973), MOE and necklace glomeruli in a cryostat. Sections were collected on Superfrost/Plus slides (Fisher Scientific) and stored at -80°C prior to further processing.

### Immunohistochemistry

Sections were first treated with EDTA buffer (1 mM EDTA, 0.05% Tween 20, pH 8.0) and microwave heating for antigen retrieval according to standard protocols (Pileri et al., 1997). Sections were then incubated in 5% normal donkey serum (NDS) (ImmunoReagents) and 2% BSA (Fisher Scientific) in 1x PBS for 1 hour at room temperature followed by overnight incubation with one or more of these primary antisera at 4°C: rabbit anti-PDE1A (H105, 1:100, Santa Cruz Biotechnology), goat anti-PDE1A (Q13, 1:100, Santa Cruz Biotechnology), goat anti-PDE2A (1:100; Santa Cruz Biotechnology), chicken anti-GFP (1:500; AVES) and rabbit anti-CART (1:1,500; Phoenix Pharmaceuticals). After washing with 1x PBS, slides were incubated in donkey anti-rabbit (1:500, Life Technologies), donkey anti-goat (1:500, Life Technologies), and donkey anti-chicken (1:500, Jackson Immuno Research) secondary antibodies for 1 hour at room temperature. No signal was obtained with each of the primary or secondary antibodies alone.

### RNAscope *in situ* hybridization

RNAscope™ *in situ* hybridization is a commercially available technology that utilizes a branched or “tree” method. We purchased *Pde1A*, *Pde2A*, *Gucy2d* and *Omp* probes from Advanced Cell Diagnostics (ACD) and used them to visualize mRNA transcripts in tissue sections of mouse olfactory epithelium according to the manufacturer’s protocol. Briefly, slides were washed in 1x phosphate buffered saline (PBS) for 5 min. Slides were then transferred to distilled water for 5-10 sec, followed by a wash in fresh 100% EtOH for 5-10 sec. Protease III (ACD) was added to each section and the slides were placed in the slide rack of a HybEZ Oven (ACD) for 30 min at 40°C. Slides were washed two times in distilled water for 5-10 sec each. All remaining incubation steps were done using this oven and tray system. Tissue sections were incubated in desired probe (~2–3 drops/section) for 2 hours at 40°C. Slides were washed four times in 1x wash buffer (ACD, 310091) for 1 min each. Amplification and detection steps were performed using the RNAscope™ Multiplex Fluorescent reagent Kit (ACD, 320851). Sections were incubated with Amp1 for 30 min at 40°C and then washed four times in wash buffer (ACD) for 1 min each. Amp2 was incubated on the sections for 15 min at 40°C, followed by four washes in wash buffer. Sections were incubated in Amp3 for 30 min at 40°C and washed four times in wash buffer for 1 min each, followed by incubation of Amp4 for 15 min at 40°C. Slides were washed four times in wash buffer for 1 min each. Sections were coverslipped with Vectashield with DAPI (Vector).

## RESULTS

Canonical olfactory sensory neurons (OSNs), which utilize cAMP as a second messenger, express multiple cyclic nucleotide phosphodiesterase isoforms (Cherry and Davis, 1995; Yan et al., 1995). We first asked if GC-D+ OSNs expressed any phosphodiesterases in addition to the cGMP-dependent phosphodiesterase Pde2a (Juilfs et al., 1997). Based on findings from a preliminary polymerase chain reaction screen of olfactory mucosa cDNA (not shown), we performed triple-label immunohistochemistry for green fluorescent protein (GFP), the known GC-D+ OSN marker Pde2a and phosphodiesterase Pde1a in GCD-EGFP reporter mice. Pde1a immunoreactivity (antisera H105) was restricted to OSNs expressing GFP and Pde2a (**Figure 1**). GGNs also express Pde2a. We found that Pde2a and Pde1a were colocalized by double-label immunohistochemistry for this chemosensory tissue (**Figure 2A**).

**Figure 1:**
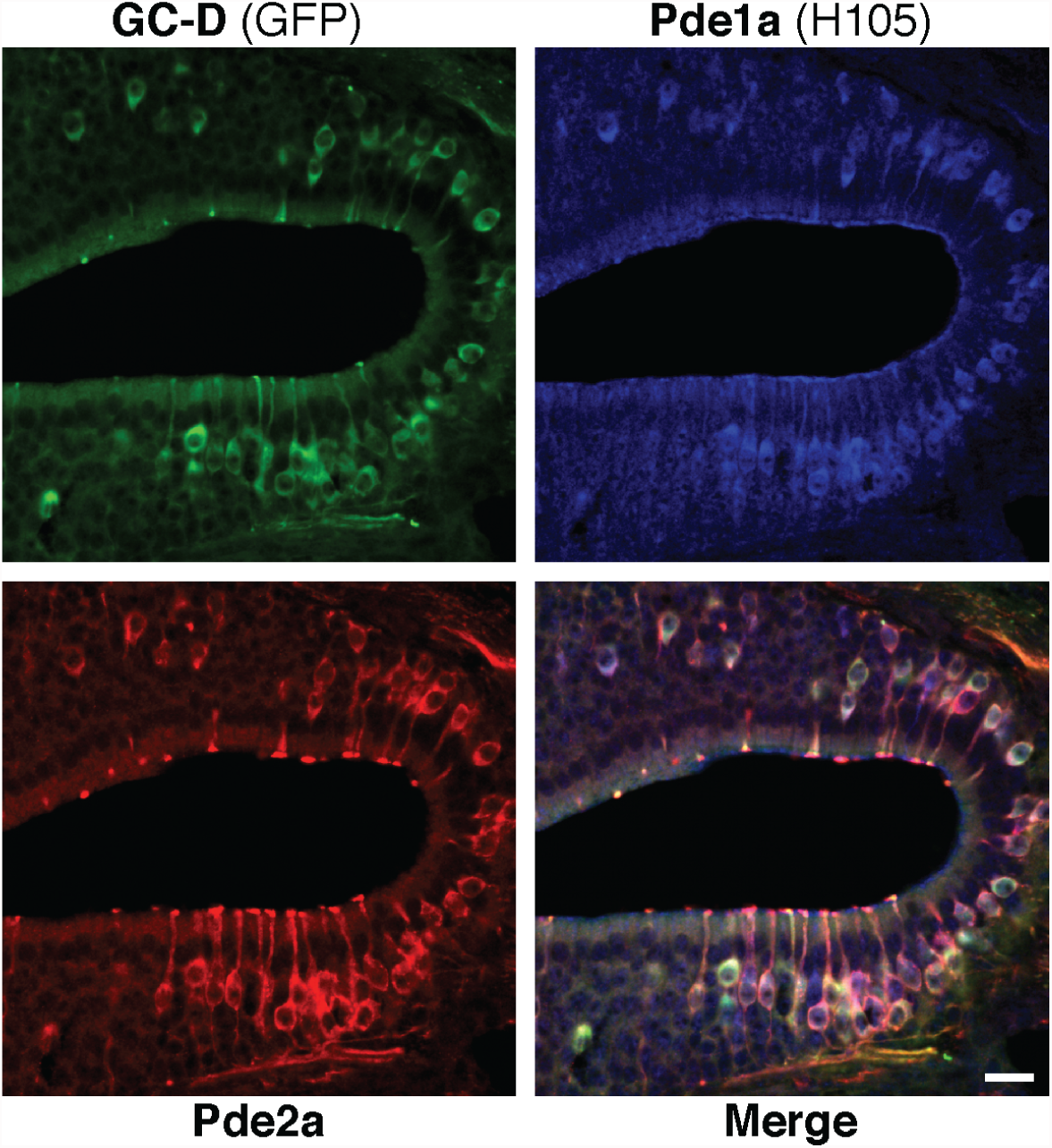
GC-D+ OSNs express two Pde isoforms. Immunohistochemical staining for GFP (green), Pde1a (blue) and Pde2a (red) in main olfactory epithelium of GCD-GFP mice. Scale bar, 20 µm.

**Figure 2:**
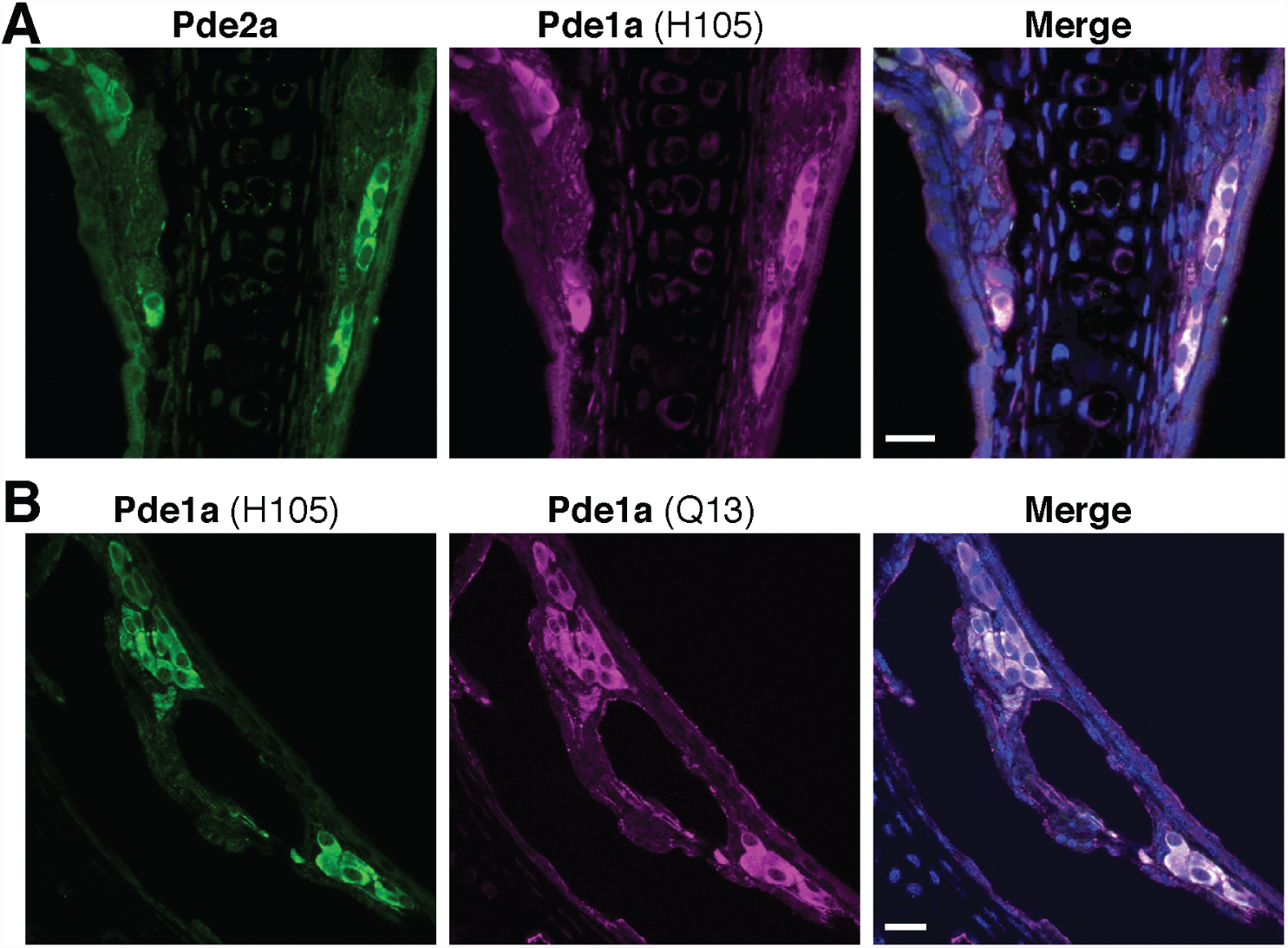
Chemosensory neurons of the Grueneberg ganglion (GGNs) co-express Pde2a and Pde1a. **(A)** Immunohistochemical staining for Pde2a (green) and Pde1a (antisera H105, magenta) colocalize in GGNs. **(B)** Pde1a antisera H105 (green) and Q13 (magenta), which target peptide antigens in different regions of the Pde1a protein and were raised in different species, colocalize in GGNs. Scale bar, 20 µm.

To confirm that the Pde1a antisera indeed recognizes Pde1a, we co-labeled the GG with two different Pde1a antisera (H105 and Q13) that were generated to distinct peptide antigens. The two immunosignals overlapped completely (**Figure 2B**). Finally, we used the *in situ* hybridization technique RNAScope™ to co-localize Pde2a and Pde1a messenger RNA in the MOE. Consistent with the immunohistochemistry results, we found that Pde2a and Pde1a message are co-localized in GC-D+ OSNs (**Figure 3**), which were visualized with another novel marker for these neurons, the CART peptide (Douglass et al., 1995). In the MOE of GCD-EGFP reporter mice, CART immunoreactivity was restricted to OSNs expressing GFP and Pde2a (**Figure 4A**). However, no CART signal was seen in GGNs (**Figure 4B**). Together, these results indicate that Pde1a is expressed by both GC-D+ OSNs and GGNs, but CART is a specific marker for GC-D+ OSNs.

**Figure 3:**
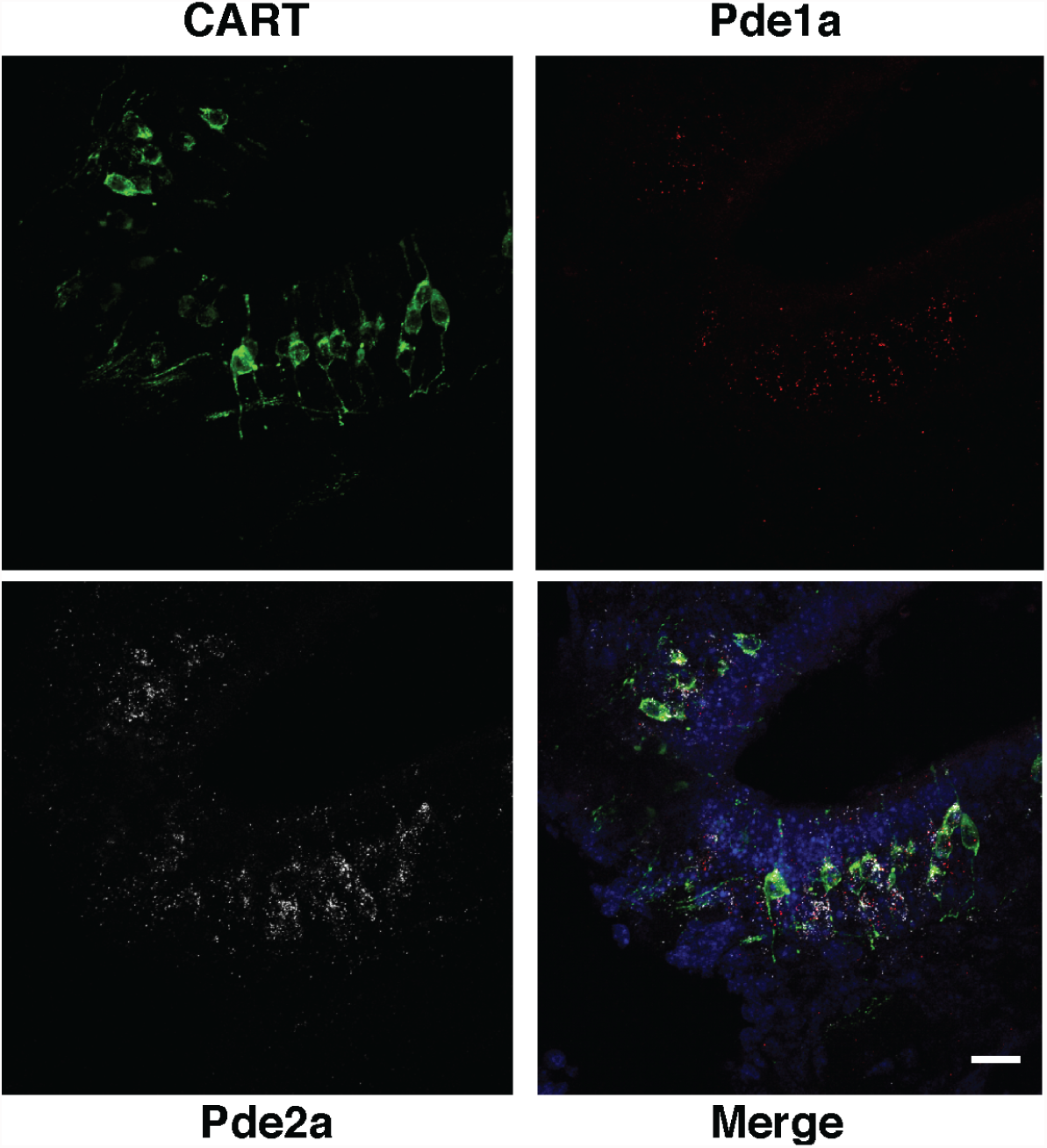
Pde1a message is co-expressed with Pde2a and CART in the main olfactory epithelium. RNAscope™ was used to visualize messenger RNA for Pde1a (red) and Pde2a (white) in tissue sections immunostained for CART (green). Scale bar, 20 µm.

**Figure 4:**
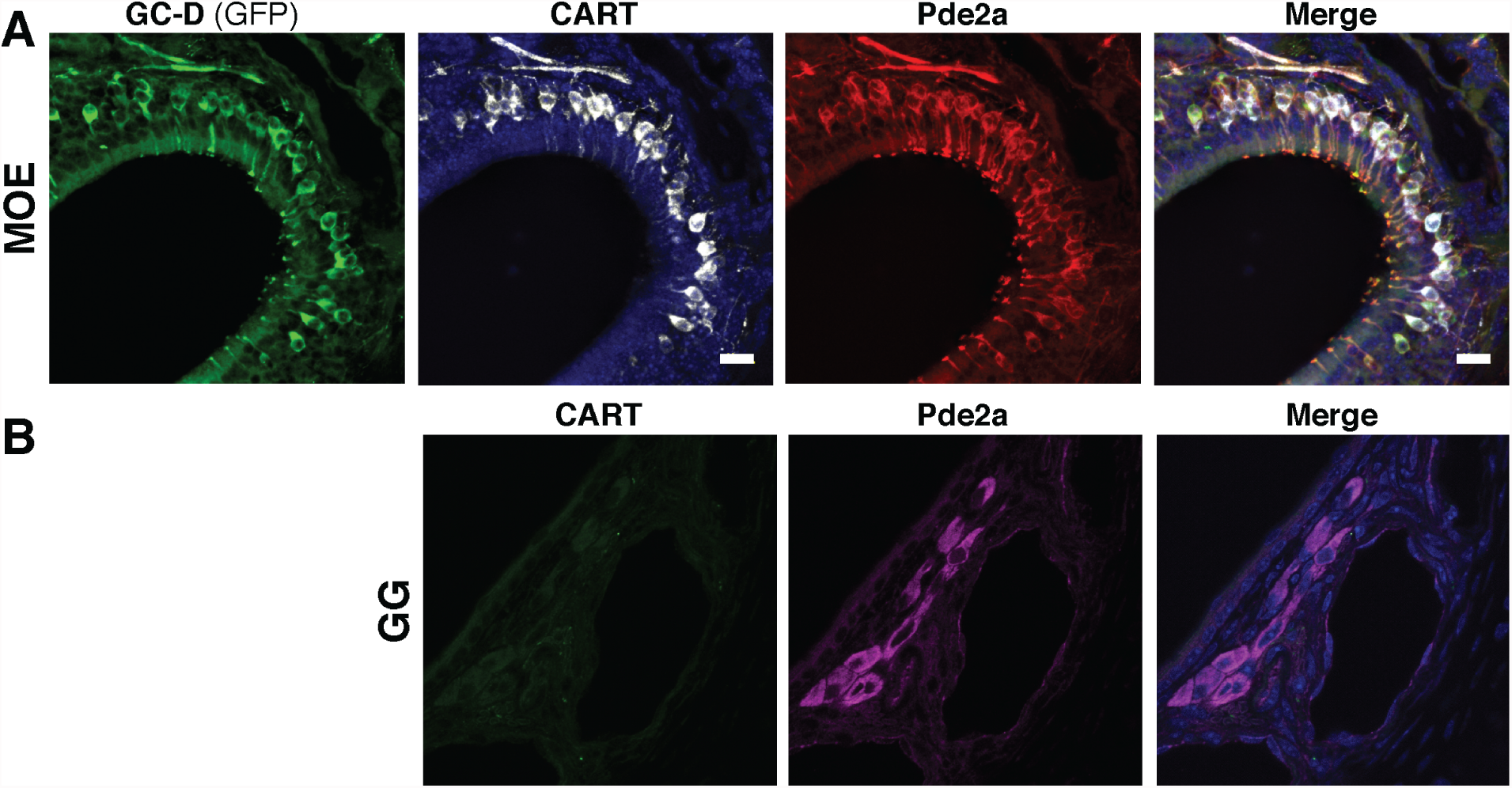
GC-D+ OSNs, but not GGNs, express CART. **(A)** Immunohistochemical staining for GFP (green), CART peptide (white) and Pde2a (red) in main olfactory epithelium of GCD-GFP mice. **(B)** No CART immunostaining (green) is seen in GGNs, which do express Pde2a (magenta). Scale bar, 20 µm.

Both GC-D+ OSNs and GGNs project to the caudal MOB where they innervate separate subpopulations of so-called “necklace” glomeruli that ring the bulb (Juilfs et al., 1997; Matsuo et al., 2012). Consistent with the results in MOE, CART immunostaining was restricted to glomeruli innervated by GC-D+ OSNs (**Figure 5**). However, the expression of the phosphodiesterases in the necklace glomeruli varied in a striking way. Pde1a immunoreactivity was very faint in GC-D-innervated necklace glomeruli, which expressed CART (**Figure 5**) or GFP (**Figure 5**). Pde2a immunoreactivity also varied between the two subsets of necklace glomeruli, with stronger Pde2a staining colocalizing with GFP-immunopositive (i.e., GC-D+) glomeruli and a weaker Pde2a signal colocalizing with strong Pde1a staining (i.e., GGN-innervated glomeruli). These results indicate that GC-D+ OSNs and GGNs (and their glomerular targets) differ in the expression of CART and in the subcellular distribution of Pde1a.

**Figure 5:**
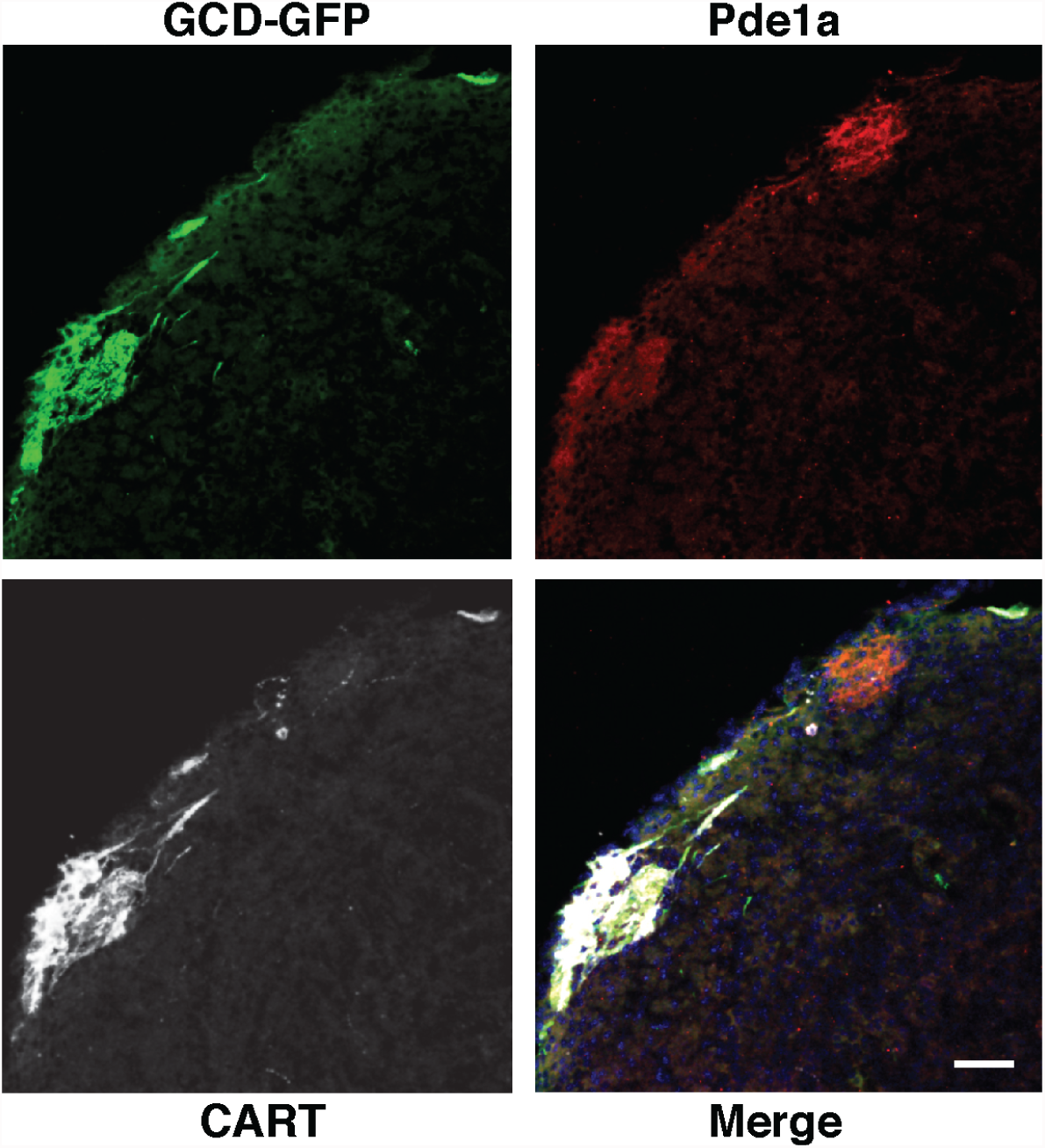
CART is expressed in necklace glomeruli innervated by GC-D+ OSNs. Immunohistochemical staining for GFP (green), Pde1a (red) and CART (white) in caudal main olfactory bulb of GCD-GFP mice. Scale bar, 20 µm.

**Figure 6:**
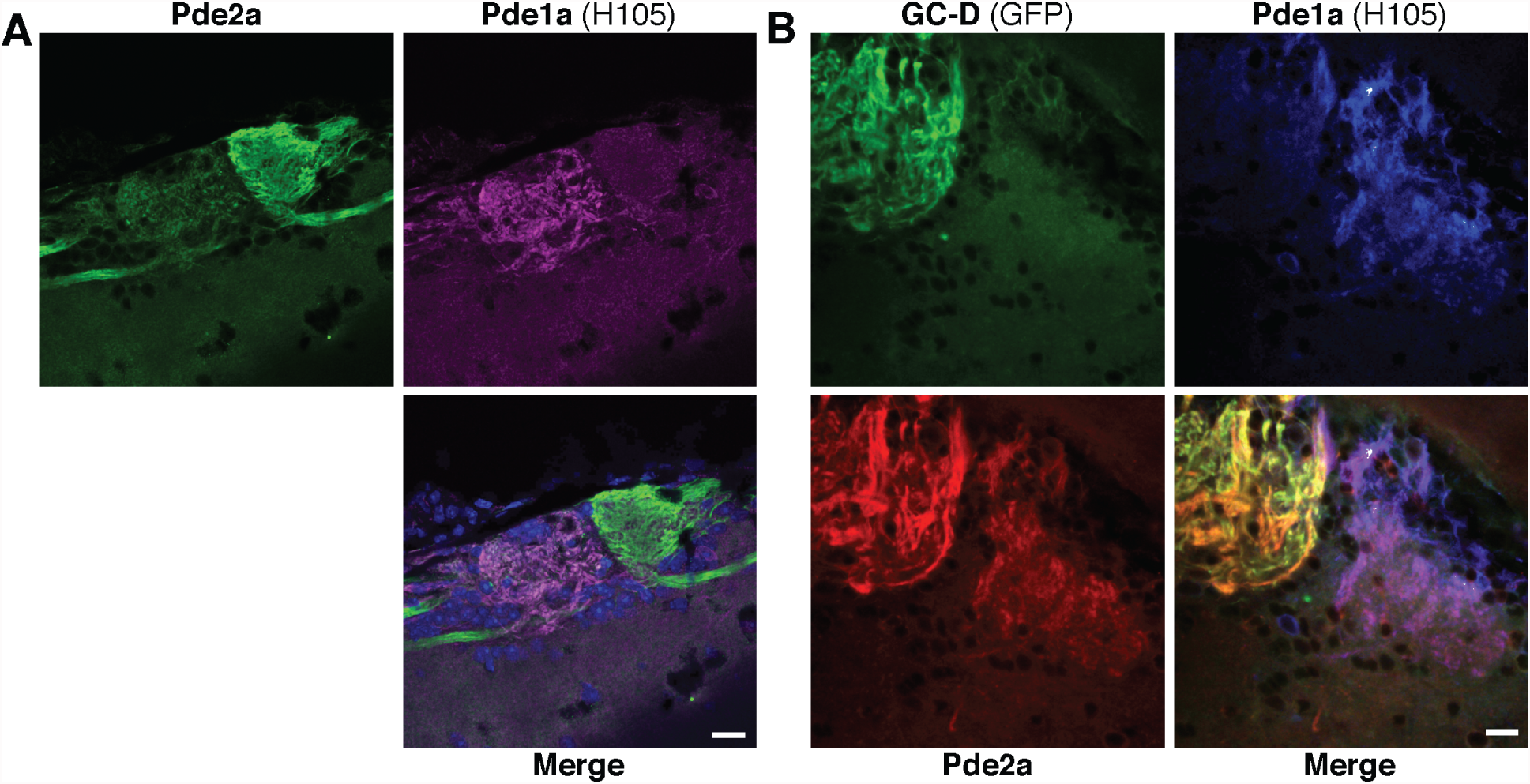
Differential expression of Pde2a and Pde1a in necklace glomeruli targeted by GC-D+ OSNs and GGNs. **(A)** Pde1a-immunopositive glomeruli (magenta) stain weakly for Pde2a (green), and vice versa. (B) Strongly Pde2a-immunopositive glomeruli (red) are innervated by GC-D+ OSNs (green), indicating that strongly Pde1aimmunopositive glomeruli (blue) are innervated by GGNs. Scale bar, 20 µm.

## DISCUSSION

GC-D+ OSNs and GGNs are morphologically distinct, anatomically distant, respond to discrete chemostimuli and mediate different olfactory behaviors. Even so, they share a similar molecular profile (**Table 1**), and their glomerular targets in the olfactory bulb are in close proximity. To facilitate functional and anatomical studies of these separate but similar olfactory subsystems, it is useful to identify new molecular markers that can be used to distinguish between or differentially target them. Here, we add two new molecular markers that expand the repertoire of tools that can be used experimentally to better understand the functions and biological roles of the olfactory subsystems defined by the GC-D+ OSNs and the GGNs.

**Table 1:**
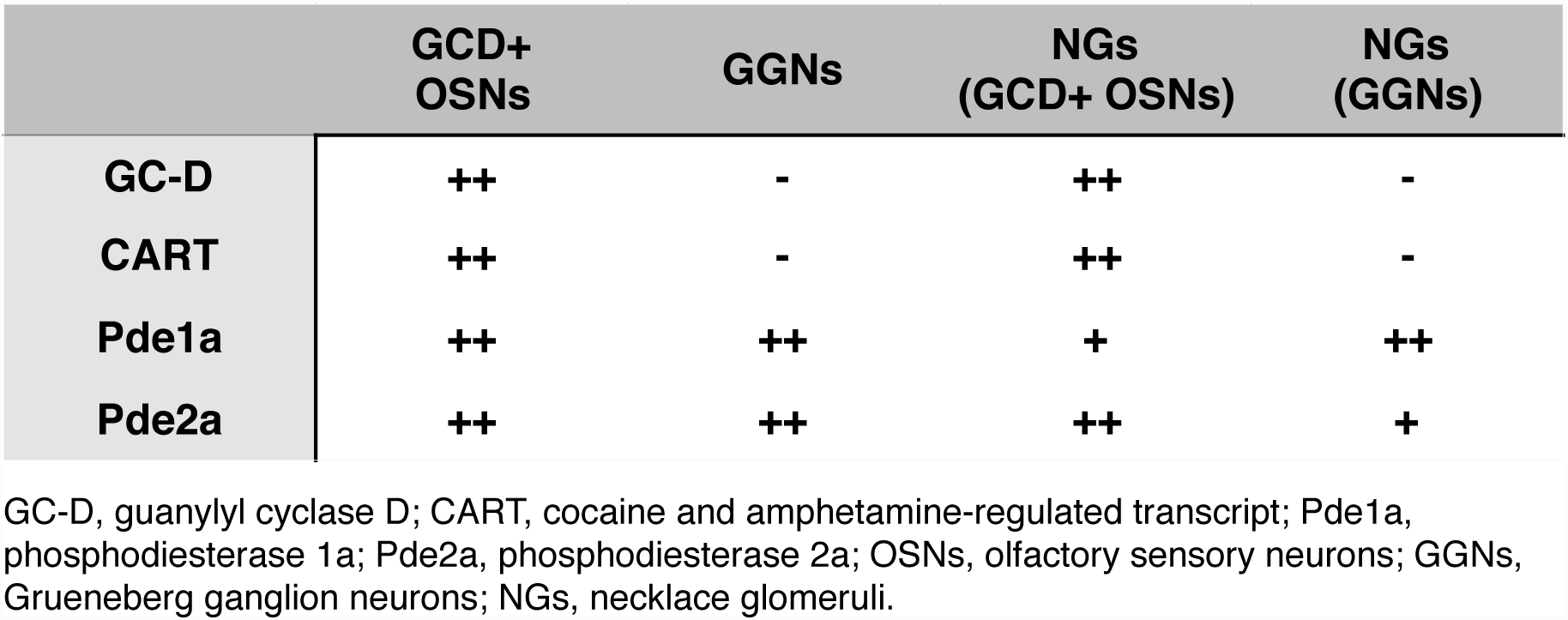
Molecular signatures of GC-D+ OSNs, GGNs, and their glomerular targets.

Pde1a is found in the cell bodies of both GC-D+ OSNs and GGNs. However, while Pde1a immunoreactivity is strong in GGN axons it is comparatively faint in GC-D+ OSN axons. This subcellular distribution is different from Pde1c, the other Pde1 isoform found in the peripheral olfactory system, which is largely restricted to the dendritic cilia of canonical OSNs (Borisy et al., 1992). Pde1a distribution in GGNs more closely resembles that of Pde4a in canonical OSNs (Cherry and Davis, 1995) or Pde2a in GC-D+ OSNs (Juilfs et al., 1997), both of which are found in the dendrites, cell bodies and axons of these cells. By contrast, Pde2a immunostaining is strong in GC-D+ OSN axons but much weaker in GGN axon terminals. The reasons for these differences in axonal, but not cell body, staining for Pde1a and Pde2a are unclear. Potential explanations include a difference in protein expression level per axon or differences in innervation density between GC-D+ OSNs and GGNs. This latter possibility seems less likely as GGN axons show stronger staining for Pde1a while GC-D+ OSN axons show stronger staining for Pde2a. Necklace glomeruli innervated by GC-D+ OSNs also receive heterogeneous afferent input, with axons from Pde2a-negative, Pde4a-negative neurons intermingling with those of the Pde2a-positive GC-D+ OSNs (Juilfs et al., 1997; Cockerham et al., 2009).

It is difficult to predict the functional role of Pde1a (or Pde2a) in either GC-DOSNs or GGNs. Pde1a and Pde2a are differently regulated (by Ca^2+^/calmodulin and cGMP, respectively), but both hydrolyze cGMP (Bender and Beavo, 2006). In the periphery, these two phosphodiesterases show similar distributions in both GC-D OSNs and GGNs. However, differences in axonal localization may reflect different roles within this cellular compartment. However, one must be careful when predicting function from location. For example, Pde1c – which is regulated by Ca^2+^/calmodulin, hydrolyzes cAMP and is enriched in the dendritic cilia of canonical OSNs – was long predicted to facilitate the termination of the odor response in these cells. Surprisingly, odor responses were terminated more rapidly in *Pde1c* null mice than in their wild type controls. Only in mice lacking both Pde1c and Pde4a (the latter which is excluded from olfactory cilia) were odor responses significantly prolonged. A better understanding of the contributions of Pde1a and Pde2a to olfactory function awaits studies in which each protein can be separately and specifically deleted or disrupted.

CART expression, like that of GC-D, clearly differentiates GC-D+ OSNs and their glomerular targets from GGNs. It’s role in these neurons remains unknown. The CART peptide is found in a number of neuronal and non-neuronal cell types (Rogge et al., 2008). It has been linked to both reward and feeding systems in the brain, but its roles in specific cell types are poorly understood. In the striatum, CART expression is upregulated in animals exposed to cocaine or amphetamine (Douglass et al., 1995). It is unclear whether CART expression is changed in GC-D+ OSNs under certain conditions, or whether it functions as a neurotransmitter or other paracrine factor for these cells. But whatever it’s role, it highlights a functional complexity in the peripheral olfactory system of mammals that remains only partially understood.

### Funding sources

National Institute on Deafness and Other Communication Disorders grants R01DC005633 and R21DC012501.

